# Surgical Protocol for a Large, Resealable Cranial Window Enabling Longitudinal, Multi-Modal Electrophysiology Recordings of Mouse Default Mode Network

**DOI:** 10.1101/2025.11.13.688186

**Authors:** Tommi Kiiso, Paula Partanen, Keyrstin Jacobs, Bendegúz Áron Varga, Satu Palva, Matias Palva, Eero Castrén, Raz Balin

## Abstract

The Default Mode Network (DMN) is a central large-scale brain network implicated in a range of cognitive functions and neuropsychiatric disorders, such as major depressive disorder (MDD). Studying the DMN’s complex dynamics in animal models provides invaluable insights into its function in both healthy and pathological states. However, performing a stable, long-term, and large-scale electrophysiological recordings from the multiple, deep, and distributed nodes of the DMN in awake, behaving mice has been a significant challenge.

Here, we present a novel, two-phase surgical protocol developed to create a large (4×7.6 mm), durable, and resealable cranial window in mice. The procedure is designed to preserve the integrity of the dura mater, which is paramount for long-term brain health and recording stability.

This window facilitates repeated, longitudinal recordings from over 1,000 electrodes simultaneously by combining surface-level micro-electrocorticography (µECoG) with two high-density intracranial electrode probes, allowing unprecedented access to the DMN. This technique provides a robust platform for multi-modal, multi-scale interrogation of network-wide electrophysiological dynamics over several weeks, opening new avenues for investigating the neuroplastic changes underlying the pathophysiology of brain disorders and for evaluating the chronic effects of novel therapeutics.

**SUMMARY:** This protocol describes a two-phase surgical method to create a large, resealable, dura-sparing cranial window in mice. This technique enables chronic, multi-modal electrophysiological recordings from distributed, deep brain networks, such as the Default Mode Network, over several weeks.

## INTRODUCTION

The Default Mode Network (DMN) was first identified in human brain imaging studies as a constellation of regions consistently suppressed during externally oriented, attention-demanding tasks^1^. It is now considered to be a fundamental large-scale network supporting internally oriented cognitive processes, such as self-referential thought, autobiographical memory, and planning of future events^2,3^. In humans, the canonical DMN comprises a set of anatomically connected core regions, including the medial prefrontal cortex (mPFC), posterior cingulate cortex (PCC), inferior parietal lobule (IPL), and the hippocampal formation^4^. Dysfunction within this network is a key feature of several neuropsychiatric disorders; in major depressive disorder (MDD), the DMN commonly exhibits hyperactivity and altered patterns of functional connectivity, which are thought to underlie core symptoms of MDD, such as rumination and negative self-focus^5,6^.

To dissect the cellular and circuit mechanisms driving DMN dysfunction, robust animal models are indispensable. A homologous DMN has been reliably identified in rodents, consisting of comparable brain regions such as the mPFC, cingulate cortex, and retrosplenial cortex (RSP), thereby validating the mouse as a translationally relevant model for mechanistic investigation^7,8^. Furthermore, chronic stress, a key factor in depression, has been shown to alter DMN connectivity in rodents in a manner similar to that observed in human MDD patients^9^.

Understanding the neuroplastic changes that occur in disease states or in response to chronic therapeutic interventions requires longitudinal observation of neural activity. A single recording session is insufficient to capture the dynamic adaptation of neural circuits over time^10^. In vivo electrophysiology offers the millisecond-scale temporal resolution necessary to study neural oscillations and synchrony, the fundamental mechanisms of network communication^11,12^. However, performing chronic, large-scale electrophysiological recordings in awake mice presents several technical hurdles. Conventional cranial windows are often too small to permit simultaneous access to the distributed nodes of a network like the DMN. Achieving stable, high-quality signals over many weeks from hundreds of channels, particularly when targeting both superficial cortical areas and deep subcortical structures, remains a major challenge.

This protocol directly addresses these limitations. Here, we demonstrate a two-phase surgical procedure that creates a large, resealable cranial window specifically designed for chronic, multi-modal electrophysiology. This method is distinguished by its combination of size, durability, and a dura-sparing technique that minimizes inflammation and glial scarring, thereby ensuring the long-term viability of the preparation. The resulting implant allows for the repeated insertion and removal of high-density recording hardware, including surface micro-electrocorticography (µECoG) grids for mesoscale local field potential (LFP) recordings and multiple Neuropixels probes for microscale, laminar-resolved single-unit and LFP activity^13,14^. This enables a true multi-scale interrogation of network dynamics over weeks, providing a powerful tool for systems neuroscience research.

## PROTOCOL

All animal experiments were approved by the national Animal Experiment Board in Finland (license number: ESACI/40845/2022) and experiments were conducted in compliance with the directive 2010/63/EU of the European Parliament regarding the protection of animal use for scientific purposes, and with the national guidelines for the care and use of laboratory animals.

### 1. Pre-operative Preparation and Anesthesia

1. Use C57BL/6JRj mice, at least 10 weeks old and weighing over 20g.
2. Administer pre-operative analgesics and anti-inflammatory agents via subcutaneous (s.c.) injection: Carprofen (5 mg/kg), Buprenorphine (0.05 mg/kg), and Dexamethasone (2 mg/kg).
3. Anesthetize the mouse with isoflurane (induction: 4%, maintenance: 1.5−2.5%). Confirm the depth of anesthesia by the loss of toe pinch reflex.
4. Secure the mouse in a stereotactic frame on a heating pad maintained at 37 °C. Apply carbomer eye ointment to prevent corneal drying (see Table of Materials).

### 2. Phase A: Headplate and Reference Implantation

1. Shave the fur from the top of the head, approximately from the eyes to the neck, and disinfect the area with povidone-iodine. Inject lidocaine-norepinephrine solution (s.c.) as a local anesthetic under the scalp skin.
2. Make a transverse incision at the ear-line and remove the skin from the top of the skull by cutting along the lateral side of the head towards the eyes, where one makes a diagonal cut towards the midline to fully expose the skull surface. Meticulously clean the skull surface with acetone to remove all periosteum, which is critical for ensuring strong adhesion of the implant.
3. Use a circular scalpel to scrape any remaining tissues and remaining deposits that were not dissolved entirely by the acetone.
4. Using the stereotactic device, mark a 4×7.6 mm rectangular area on the skull relative to bregma. The rectangle should extend 2 mm laterally from bregma on both sides, 3 mm rostrally, and 4.6 mm caudally.
5. Mark the two insertion sites for the Neuropixels probes on the right hemisphere.
  1. Mark the rostral probe site at 1.66 mm rostral and 1.95 mm lateral from bregma.
  2. Mark the caudal probe site at 2.2 mm caudal and 1.9 mm lateral from bregma.
6. Using a #11 surgical blade, engrave a crisscross pattern on all exposed bone surfaces *outside* the designated window area to enhance the grip of the adhesive and dental cement.
7. Apply a thin, even layer of cyanoacrylate glue to the engraved skull surface. Allow the glue to dry for 7 min.
8. Drill a small pilot hole over the left cerebellum, make sure to avoid penetrating the dura with the drill. Insert a reference socket in the hole using cyanoacrylate glue and reinforce with permanent dental cement. Cure the dental cement with a UV light for 1 minute.
9. Coat the bottom and top of the metal headplate with a thin layer of cyanoacrylate glue and affix it to the skull, ensuring it is centered and level.
10. Reinforce the structure by applying dental cement around the base of the headplate to create a robust, sealed enclosure. Cure the dental cement with a UV light.
11. Allow the mouse to recover from anesthesia. Return the mouse to its home cage for at least 48 h. **NOTE:** This 48 h recovery period is critical. It allows the animal to recover from the initial physiological stress and inflammation before the craniotomy, which significantly improves surgical success rates and the long-term health of the implant.

### 3. Phase B: Chronic Cranial Window Creation

1. Re-anesthetize the mouse with isoflurane (induction 4%, maintenance 1.5−2.5%), administering the same medications as in step 1.2, with the exception of the local anesthetic. Position the mouse in the stereotactic frame as in step 1.4.
2. Using a dental drill, begin thinning the bone carefully along the rectangle markings marked in phase A (section 2.4). Regularly apply ice-cold sterile artificial cerebrospinal fluid (ACSF) to the area to prevent thermal damage to the underlying cortex and to minimize bleeding.
3. Drill shallow grooves outside of the cranial window at the marked probes insertion coordinates (described in section 2.5) to serve as guides for later electrophysiological recordings.
4. **CRITICAL STEP:** Continue drilling along the rectangle until the bone is approximately 90% thinned. The bone plate should appear translucent and flexible. *Do not drill completely through the skull*.
5. **CRITICAL STEP:** Switch to a fine dura hook. Carefully insert the tip under the edge of the thinned bone and gently move it along the perimeter to detach the bone plate from the dura mater and surrounding skull. This technique is key to avoiding dural tears and preserving the long-term health of the underlying brain tissue.
6. Carefully lift the bone plate to expose the brain. Gently clear the exposed area of any coagulated blood with repeated washes of ice-cold ACSF. The dura should appear intact and transparent. **NOTE:** Be extremely cautious when lifting the bone fragment to do so in a slow and gradual manner to minimize the risk of injury to the venous sinuses, on the brain surface when lifting the bone plate.
7. Once the bleeding subsided, cut a sterile polydimethylsiloxane (PDMS) thin (<1 mm) membrane to the size of the window opening and place it directly onto the dural surface. Using a pointy tip of non-fibrous tissue to dry the remaining ACSF solution and help adhere the PDMS membrane to the dura surface. **NOTE:** The PDMS membrane protects the brain surface and applies gentle pressure, mimicking natural physiological conditions.
8. Seal the craniotomy by applying a biocompatible elastomer sealant around the edges, creating a complete, yet removable, seal around the PDMS sheet.
9. To protect the window during housing, affix a custom 3D-printed plastic cap onto the headplate using a cyanoacrylate glue.

### 4. Post-Operative Care and Habituation

1. Transfer the mouse to a clean cage for individual housing to prevent damage to the implant.
2. Monitor the mouse health closely for at least 2 days post-surgery. Administer additional analgesics (s.c.) for two days post-surgery if the mouse shows signs of pain based on the mouse grimace scale^15^.
3. Allow the mouse to recover for at least 4 days. Then, begin a 5-day habituation period of increasing duration starting with 5 min on the first day, followed by 15 min, 30 min, 45 min, and 60 min on the second, third, fourth, and fifth day, respectfully to accustom the mouse to handling and the electrophysiological recording platform.

### 5. Procedure for Longitudinal Electrophysiological Recordings

1. On a recording day, head-fix the awake mouse on the recording platform by holding the mouse metal head implant and guiding him slowly towards the head-fixation clamps.
2. Using a circular scalpel, carefully remove the protective cap and gently peel away the silicone sealant. Remove the PDMS membrane to expose the dura mater.
3. Place the µECoG grids on the cortical surface, targeting the anterior (ACC) and posterior (RSP) DMN regions.
4. Connect the ECoG external reference wire to the designated reference socket.
5. Using micromanipulators, slowly position the two Neuropixels probes close to the designated insertion points described in section 2.5 and marked on the skull in the craniotomy surgery section 3.3 adjacent to the insertion points to determine the precise location for dural puncture. location for dural puncture. Create a dura puncture device by affixing sterile 30G needle on top of a cotton swab, then bent the tip of the needle 90° with fine tweezers.
6. Once taking note of the precise location for puncture, retract both probes slightly to protect the brittle electrode from unintended damage caused by the puncturing maneuver and carefully puncture the dura mater without damaging the brain tissue.
7. Using micromanipulators, slowly insert the two Neuropixels probes through the dura at the pre-marked coordinates, both at depth of 5 mm, to target the DMN regions of interest. **NOTE:** The small dural hole created by the 30G needle will be plugged within several minutes, therefore it is of outmost importance to insert the probe shortly after the dural puncture was made.
8. Begin data acquisition using a suitable data acquisition system (e.g., OpenEphys).
9. After the recording session, gently retract the probes and remove the grids. Replace the PDMS membrane with a new one and apply a fresh layer of elastomer sealant and the protective cap. Return the mouse to its home cage. This process can be repeated for multiple recording sessions over several weeks.

## REPRESENTATIVE RESULTS

A successful surgery results in a clear, transparent window over the cortex, with visible vasculature and minimal signs of inflammation or infection. This clarity can be maintained for over 21 days (Figure 1), enabling true longitudinal studies. The robustness of the implant is demonstrated by its ability to remain stable even throughout a 21-day chronic corticosterone administration paradigm, a common model for inducing depression-like states in rodents^10,16,17^. This confirms the suitability of the method for use in chronic disease modeling.

**Figure 1.**
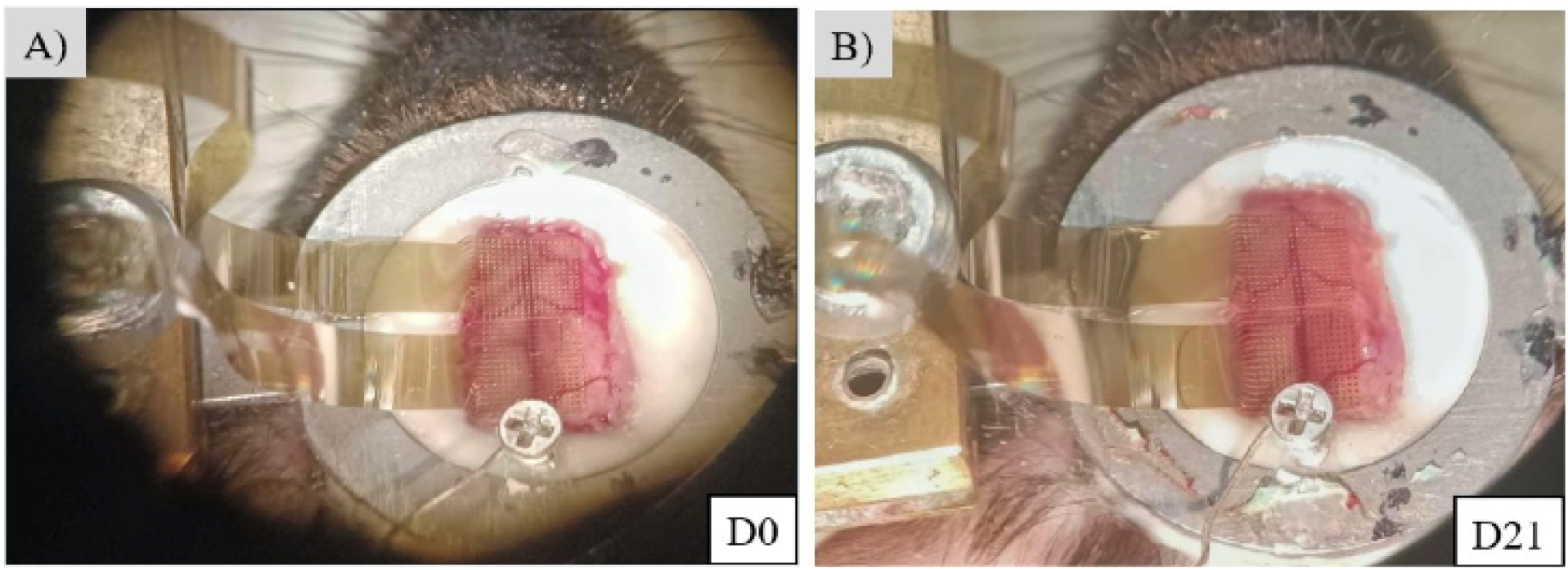
Representative images of a chronic cranial window over 21 days. The dura mater remains clear and the underlying vasculature healthy, with minimal signs of inflammation or gliosis. This stability is a direct result of the dura sparing technique and is essential for longitudinal studies. A) Day 0, B) Day 21.

The primary validation of this technique is the acquisition of stable, high-quality, multi-modal electrophysiological data over time. The resealable window allows for the repeated insertion of probes to record from the same neuronal populations across weeks. As shown in Figure 2, stable single-unit waveforms can be isolated from the same Neuropixels channel during a baseline session and again three weeks later, demonstrating the precision and low-trauma nature of the re-insertion process (Figure 2). This stability is the foundation for performing reliable, advanced analyses of network dynamics, such as phase-locking value (PLV) and dynamic functional connectivity (DFC), across chronic experimental timelines.

**Figure 2.**
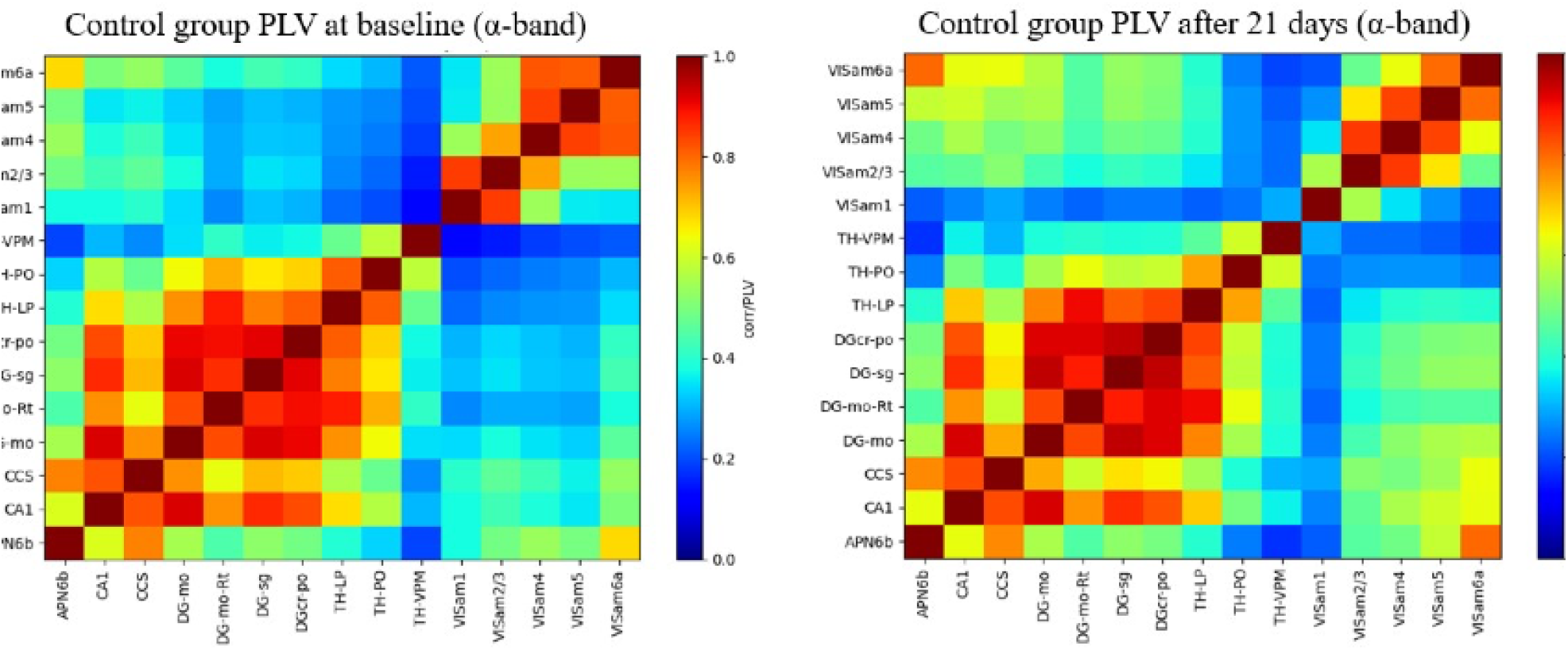
Representative α-band (8-12 Hz) phase locking value matrices from caudal intracranial electrode probe. Probes were targeting caudal regions of the DMN including visual area layers (VISam), thalamic (TH) and hippocampal areas (DG, CA1), and posterior parietal association area (APN6b). (n = 3)

To further validate the technique, immunohistochemical stainings were performed for GFAP and IBA1 in post-mortem brain slices over three weeks after the craniotomy surgery. GFAP are expressed by astrocytes which are upregulated in reactive gliosis such as brain injury or inflammation. IBA1 on the other hand is expressed by microglia which are more active during neuroinflammatory processes. We see no significant differences in either of the markers between no-surgery control group and vehicle group that underwent surgery (Figure 3).

**Figure 3.**
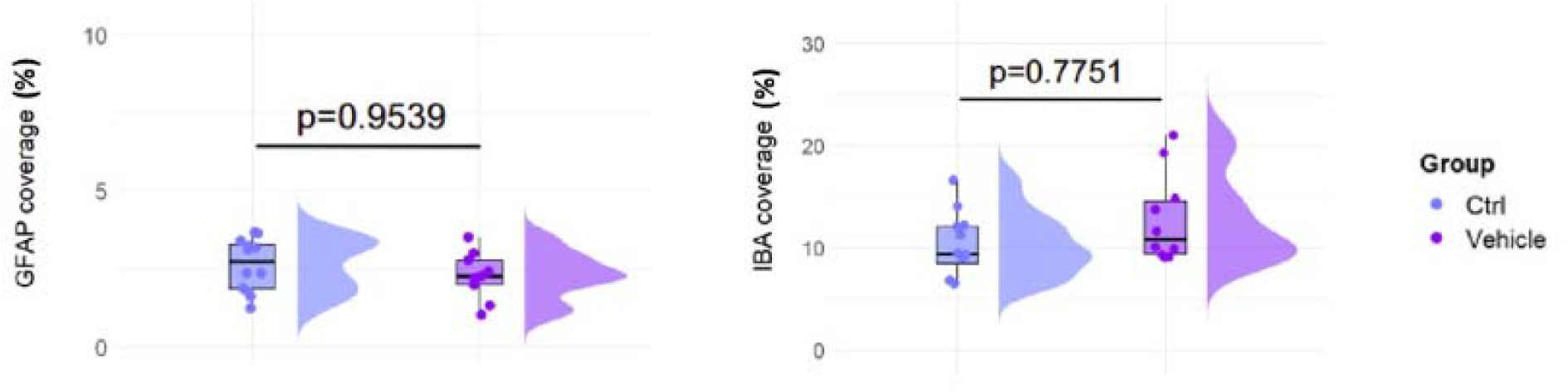
Immunohistochemistry analysis for GFAP (astrocytes) and IBA1 (microglia) from no-surgery control group, and vehicle group that underwent surgery. In the expression of GFAP and IBA1, there is no significant differences between control and vehicle groups, suggesting that the surgery does not result in significant inflammation or reactive gliosis in the brain. Each data point represents one brain slice (n = 3)

## DISCUSSION

This report details a two-phase, dura-sparing surgical protocol that provides a reliable and highly effective method for creating a large, chronic, and resealable cranial window in mice. The significance of this technique extends beyond the size of the window; its principal advantage is the capacity for simultaneous, multi-scale interrogation of large-scale brain networks over longitudinal timescales. By combining surface µECoG grids (providing mesoscale LFP data) with deep, high-density Neuropixels probes (providing microscale, laminar-resolved single-unit and LFP data), researchers can directly investigate how cell-type-specific activity within deep DMN nodes contributes to the emergent, network-level dynamics observed on the cortex. This approach facilitates a move from purely correlational observations toward a mechanistic dissection of network function.

Two aspects of the protocol are fundamental to its success. First, the division of the surgery into two phases separated by a 48-hour recovery period is a critical design choice. This strategy minimizes the cumulative physiological stress on the animal, reduces the acute inflammatory response, and allows for initial healing before the more invasive craniotomy. This significantly improves the probability of a successful, healthy long-term implant. Second, the dura-sparing craniotomy technique is the single most critical maneuver. By thinning the bone to approximately 90% of its thickness and then gently lifting the bone flap with a hook, the integrity of the dura mater is preserved. The dura acts as a vital protective barrier; keeping it intact is essential for preventing the inflammation, glial scarring, and infection that are the primary causes of failure in most chronic window preparations. This step is what enables the “resealable” and truly chronic nature of our window, permitting repeated access to the brain over many weeks with minimal damage.

The presented technique establishes a robust platform for addressing fundamental questions in systems neuroscience that require longitudinal observation. It is particularly well-suited for studying the effects of chronic stress on network plasticity and for modeling neuropsychiatric disorders like depression^10,18,19^. A key future application is the evaluation of the network-level impact of novel, long-acting therapeutics. For instance, recent work has shown that antidepressants and psychedelics can induce a state of “juvenile-like” plasticity by directly binding to the TrkB receptor^20^. This chronic window preparation is an ideal system for tracking how such compounds rewire DMN circuits over the weeks required for their therapeutic effects to manifest.

The primary limitations of this method are its technical demands. The surgery requires significant practice and a high degree of microsurgical skill to perform reliably. Furthermore, despite the use of precise stereotactic coordinates, minor variability in probe placement between recording sessions is unavoidable. For studies where the precise anatomical location is paramount, post-mortem histological verification of probe tracks is strongly recommended to confirm targeting. Finally, the current sealed-window design does not readily accommodate local pharmacological manipulations via cannula, although modifications to the 3D-printed cap and sealant procedure could be explored to incorporate such access ports.

Potential complications are manageable. Mild brain swelling may occur, particularly following a chronic treatment period, but this does not typically preclude high-quality recordings and can be managed with careful monitoring. While precise stereotactic measurements are crucial, minor variability in probe and grid placement is a potential issue. Therefore, for ultimate confirmation of recording sites, probe trajectories can be verified post-mortem using standard histological tissue staining techniques.

In conclusion, this two-phase surgical protocol overcomes major obstacles in chronic, large-scale electrophysiology. By enabling stable, multi-week, multi-modal recordings from distributed nodes of the DMN, this method provides a powerful platform to link circuit dynamics to complex behavior and pathology, ultimately advancing our ability to understand and treat complex brain disorders.

## DISCLOSURES

EC is a co-founder, Board member and recipient of research support from Kasvu Therapeutics, Ltd, and has received a speaker fee from Janssen-Cilag. The other authors have nothing to disclose.

## ACKNOWLEDGEMENTS

This work was supported by the Research Council of Finland grants #327192 and 347358, and the Sigrid Jusélius foundation. The authors thank the members of the Castrén and Palva laboratories for their technical assistance and helpful discussions throughout this project.

## MATERIALS

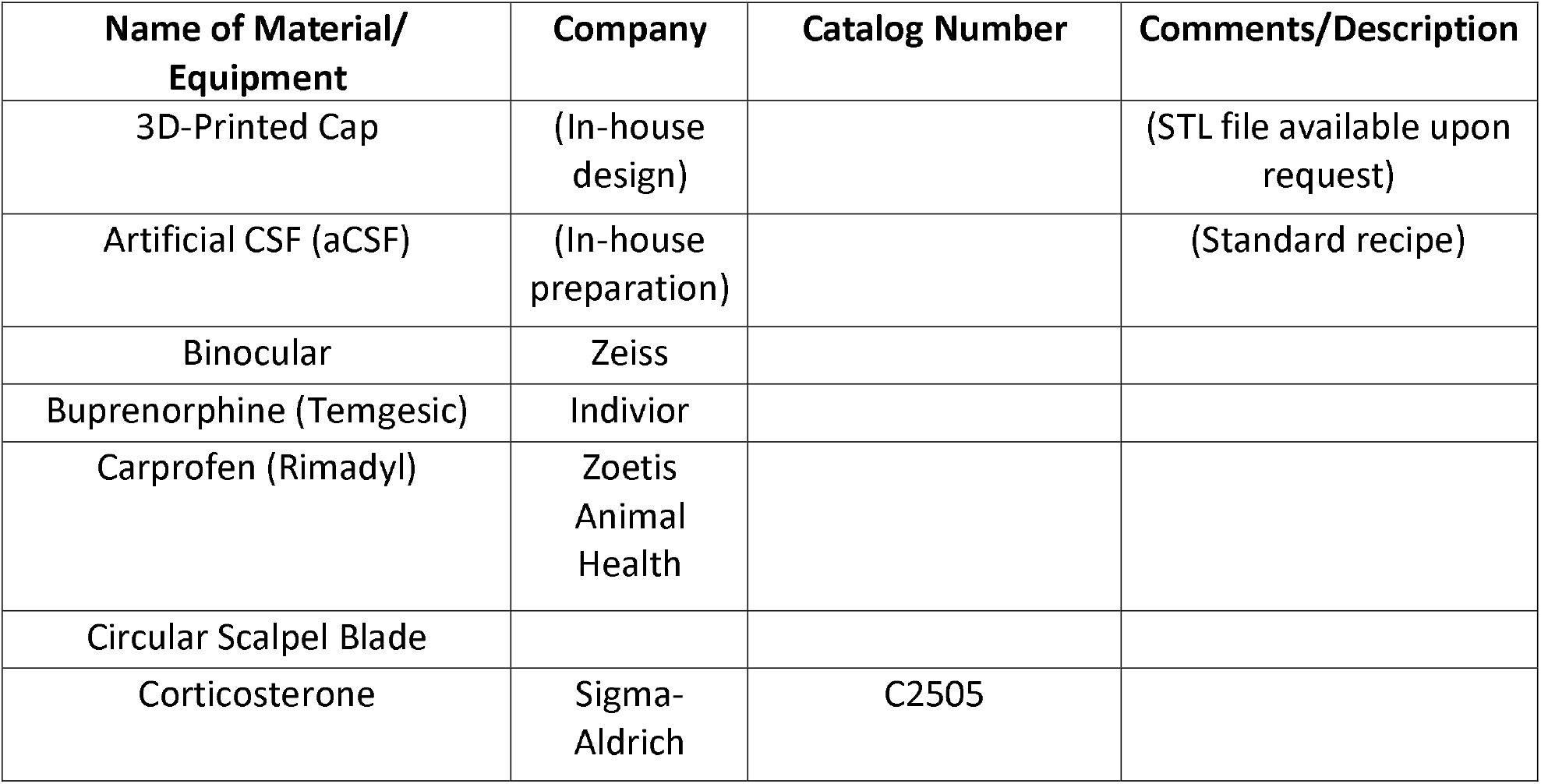
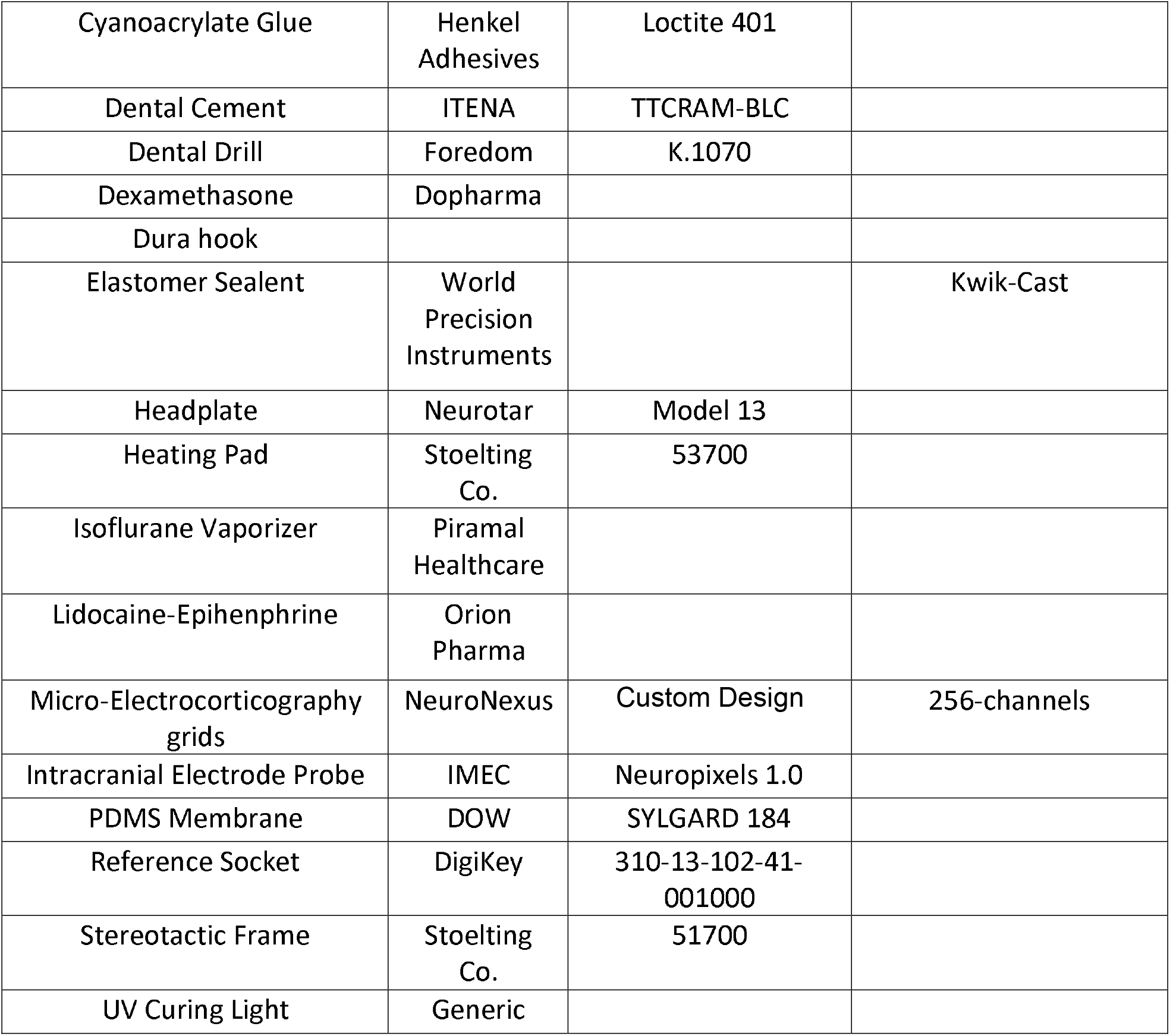

| Name of Material/<br>Equipment | Company | Catalog Number | Comments/Description |
| --- | --- | --- | --- |
| 3D-Printed Cap | (In-house design) |  | (STL file available upon request) |
| Artificial CSF (aCSF) | (In-house preparation) |  | (Standard recipe) |
| Binocular | Zeiss |  |  |
| Buprenorphine (Temgesic) | Indivior |  |  |
| Carprofen (Rimadyl) | Zoetis Animal Health |  |  |
| Circular Scalpel Blade |  |  |  |
| Corticosterone | Sigma-Aldrich | C2505 |  |
| Cyanoacrylate Glue | Henkel Adhesives | Loctite 401 |  |
| Dental Cement | ITENA | TTCRAM-BLC |  |
| Dental Drill | Foredom | K.1070 |  |
| Dexamethasone | Dopharma |  |  |
| Dura hook |  |  |  |
| Elastomer Sealent | World Precision Instruments |  | Kwik-Cast |
| Headplate | Neurotar | Model 13 |  |
| Heating Pad | Stoelting Co. | 53700 |  |
| Isoflurane Vaporizer | Piramal Healthcare |  |  |
| Lidocaine-Epihenphrine | Orion Pharma |  |  |
| Micro-Electrocorticography grids | NeuroNexus | Custom Design | 256-channels |
| Intracranial Electrode Probe | IMEC | Neuropixels 1.0 |  |
| PDMS Membrane | DOW | SYLGARD 184 |  |
| Reference Socket | DigiKey | 310-13-102-41-001000 |  |
| Stereotactic Frame | Stoelting Co. | 51700 |  |
| UV Curing Light | Generic |  |  |

## Notes

https://helsinkifi-my.sharepoint.com/:v:/g/personal/kiitommi_ad_helsinki_fi/IQCnnVkTe2PbQL-BNktg0w2qAcklbUukST8buqyRBmZMVQA?e=uQHt0z

### Summary of Updates

Formatting updated and figure positions corrected.

